# Preferential assimilation, metabolism, and transfer of organic nitrogen to host plants by Mucoromycotina ‘fine root endophytes’

**DOI:** 10.1101/2024.07.31.606011

**Authors:** Nathan OA Howard, Alex Williams, Emily Durant, Silvia Pressel, Tim J Daniell, Katie J Field

## Abstract

- Mucoromycotina ‘fine root endophytes’ (MFRE) are an understudied group of plant fungal symbionts that usually co-occur with arbuscular mycorrhizal fungi. The functional significance of MFRE in plant nutrition remains under-explored, particularly their role in plant N assimilation from the variety of sources typically found in soils.
- Using four ^15^N-labelled N sources to track N transfer between MFRE and *Plantago lanceolata,* applied singly and in tandem, we investigated N source discrimination, preference, and transfer to host plants by MFRE. We traced movement of ^14^C from plants to MFRE to determine the impact of N source type on plant C allocation to MFRE.
- We found MFRE preferentially transferred N derived from glycine and ammonium to plant hosts over that derived from nitrate and urea, regardless of other N sources present. MFRE mycelium supplied with glycine and ammonium contained more plant-derived carbon than those supplied with other N sources.
- We show that MFRE directly assimilates and metabolises organic compounds, retaining C to meet its own metabolic requirements and transferring N to plant hosts. Our findings highlight diversity in function of endomycorrhizal associations with potentially profound implications for our understanding of the physiology and ecology of plant-fungal symbioses.

## Introduction

Soils are dynamic environments where moisture, temperature, pH, and nutrient balance all vary over geographical gradients and time. Nitrogen (N), a major plant nutrient, is usually present in soil in many forms simultaneously, both organic and inorganic and at variable concentrations (Matsumoto *et al*., 2000). The majority of soil N is bound within complex, organic molecules derived from the decay of plant, animal, and microbial matter (Bremner 1949, Stevenson, 1994; Greenfield, 2001) while inorganic N, including plant-accessible ammonium and nitrate salts (Tischner, 2000, Matsumoto et *al*., 2000), accounts for a much smaller pool with high turnover rates (Jackson *et al*., 1988). The form and abundance of N in soils can be affected by both natural and anthropogenic factors, including atmospheric deposition (Chen *et al*., 2018; Moore *et al*., 2020), direct application of agricultural fertilisers (Suzuki *et al*., 2017), and through natural processes such as death and decay (Greenfield, 2001; Keenan *et al*., 2023). The resultant heterogeneity has repercussions for many biotic processes, from the production of microbial N-degrading enzymes (Fujita *et al*., 2018) to the symbioses formed between plants and mycorrhizal fungi.

Mutualistic mycorrhizas are formed between most plants and certain groups of soil fungi, across nearly all habitats on Earth (Smith and Read, 2008). These interkingdom partnerships enhance plant host acquisition of soil N and phosphorus (P), while the fungal partners benefit through provision of carbon-rich derivatives of photosynthesis, including sugars and lipids (Shachar-Hill *et al*., 1995, Keymer *et al*., 2017). The most widespread type of mycorrhizal symbioses are those formed between ∼72% of plants and arbuscular mycorrhizal (AM) fungi belonging to the subphylum Glomeromycotina (Brundrett and Tedersoo, 2018). AM fungi play an important role in supplying plant hosts with N, particularly in environments where N may be limiting (Hodge and Storer, 2015). Soil N concentration affect the frequency of root colonisation and efficiency of nutrient transfer in mycorrhizal relationships (Johnson *et al*., 2005, Bonneau *et al*., 2013) with greater N availability in soils affecting the amount of AM fungal colonisation of roots (Johnson *et al*., 2005; Solaiman *et al*., 2010; Bonneau *et al*. 2013) and N limitation driving increased AM-mediated plant N assimilation from soil (Johnson *et al*., 2010). As such, soil fertility, including N availability, is a key environmental control on AM benefits derived by host plants by influencing plant C for nutrient (N and P) exchange and host plant photosynthetic capacity (Johnson *et al*., 2015; Thirkell et al., 2016).

While research on endomycorrhizas has been dominated by AM fungi for decades, another group of soil fungi are emerging as physiologically, ecologically, and evolutionarily important root endophytes (Hoysted *et al*., 2018, Howard *et al*., 2022; Liu *et al*., 2024). Recent advances in molecular detection methods (Bidartondo et al., 2011) have revealed that members of the fungal subphylum Mucoromycotina often colonise plant roots and other tissues (e.g. nonvascular plant thalli) of various plant species, often in co-colonisation with AM fungi (Rimington *et al*., 2015, Field *et al*., 2016). Mucoromycotina is a large subphylum, sister to Glomeromycotina (Schüßler and Walker, 2011), and consists mainly of saprotrophic and pathogenic species (Hoffmann *et al*., 2013). Mycorrhizal fungi within this subphylum appear to be restricted to the order Endogonales (Hoysted *et al*., 2023, Hoysted *et al*., 2019, Hoysted *et al*., 2021b) and are referred to as Mucoromycotina ‘fine root endophytes’ (MFRE). Like AM fungi, MFRE form associations with a wide range of host plants (Kowal *et al*., 2020; Albornoz *et al*., 2021) across diverse habitats, from western Europe (Field *et al*., 2015) to Australasia (Albornoz *et al*., 2022), each with varying edaphic factors including moisture (Deng *et al*., 2020), organic matter (Stockmann *et al*., 2015) and nutrient status (Zhang *et al*., 2019). Whilst recent investigations have described some key aspects of MFRE function and biology (Orchard et al., 2017a; Field et al., 2019; Hoysted et al., 2021a, Hoysted et al., 2023, Hoysted et al., 2019; Albornoz et al., 2021a; Hoysted *et al*., 2023) the contributions of MFRE to host plant nutrition and responsiveness to environmental factors remain relatively poorly understood, rendering insights into the significance of MFRE in plant communities, soil ecology and nutrient cycling unclear. Establishing how the availability of nutrients impact the MFRE-plant symbiosis is an important first step in determining their significance in both natural and agricultural ecosystems.

Co-colonisation of plants by both MFRE and AM fungi occurs frequently in natural habitats (Ryan and Kirkegaard, 2012; Orchard *et al*., 2017b). As such, there is strong potential for functional complementarity between the two fungal groups. In the liverworts *Allisonia* and *Neohodgsonia*, dual MFRE-AM fungal associations appear to be functionally complementary in terms of supplying host plants with N and P (Field *et al*., 2016). Vascular plants also gain nutritional benefits directly from associations with MFRE in terms of fungal transfer of ammonium-N and P to host plants in return for plant-fixed C resources (Hoysted *et al*., 2019; 2021; 2023), even in the absence of other microbes within monoxenic microcosms (Hoysted *et al*., 2023). Experimental evidence for MFRE involvement in vascular plant N acquisition is currently limited to a single source of inorganic N, ammonium chloride (Hoysted *et al*., 2023) and it is unknown how N availability within the environment influences MFRE function. Ammonium (NH_4_^+^)-N is preferentially transferred to host plants by AM fungi over other N-containing compounds (Johansen *et al*., 1996; Toussaint *et al*., 2004) and is therefore often used in experiments (e.g. Ames *et al*., 1983; Yang *et al*., 2014). Considering the putative saprotrophic capabilities of MFRE (Field *et al*., 2015, 2019; Hoysted *et al*., 2023) and the recent indication that these fungi provide N derived from complex organic matter to liverwort hosts (Field *et al*., 2019), it is probable that MFRE access, assimilate and transfer N from a variety of sources in the soil. Given what is known from AM symbioses, it seems likely that the availability of N within the substrate has an impact on the quantity of N transfer by MFRE to host plants, with greater plant-available N concentrations driving lower rates of transfer of N from any source to the host plant.

Inorganic sources of N, such as NH_4_^+^ and nitrate (NO_3_^-^), are simple molecules and offer a relatively low energetic cost of assimilation by symbiotic fungi compared to more complex organic compounds containing N. In AM fungi, NH_4_^+^ is preferentially assimilated over NO_3_^-^, likely because of the higher energetic cost associated with NO_3_^-^ reduction (Johansen *et al*., 1996; Marzluf, 1997; Toussaint *et al*., 2004). The energetic cost of N assimilation by AM fungi is met through supply of hexoses and lipids by the host plant (Shachar-Hill *et al*., 1995; Keymer *et al*., 2017). Given that MFRE are facultative saprotrophs (Field *et al*., 2015; Hoysted *et al*., 2023), it is possible that at least some of the energetic cost of assimilation of N from the soil may be ameliorated through saprotrophic C acquisition. This would provide MFRE with a physiological niche distinct from AM fungi, offsetting MFRE demand on host plant C resources by assimilation of soil C, while providing plants with access to nutrients from a wider pool of sources in the soil. Despite this, it is possible that MFRE preferentially assimilate inorganic N due to the extra carbon cost of metabolising N of organic origin, as is the case for AM fungi (Johansen *et al*., 1996; Marzluf, 1997; Toussaint *et al*., 2004). However, given that MFRE is not obligately biotrophic (Field *et al*., 2015), the case may be that organic compounds represent an important source of necessary carbon for MFRE.

Using *Plantago lanceolata*, a common host for both AM and MFRE fungi, colonised by an MFRE isolate (*Lyc-1*; Hoysted *et al*., 2019; 2023) in controlled monoxenic microcosms, we investigated the ability of MFRE to access, assimilate, and transfer ^15^N from a selection of inorganic and organic compounds commonly found in soils. With a series of single ‘N source’ and ‘fungal choice’ experiments, we simultaneously quantified the allocation of host plant photosynthates are passed to MFRE mycelium and fungal acquired N transfered to the host across multiple N sources and availabilities, and consider possible underpinning mechanisms.

## Materials and Methods

### Fungal inoculum

MFRE isolate *Lyc-1,* initially isolated from *Lycopodiella inundata* (Hoysted *et al*., 2019; 2023), was maintained on Gamborg B5 basal medium at 50% concentration (1.6g.L^-1^; Sigma-Aldrich; 187.4 µg.g^-1^N; table S3) buffered with 0.5g.L^-1^ MES (Sigma-Aldrich) solidified with 1% agar (referred to as ½GB5). Cultures were kept in the dark and incubated at 25°C. Immediately following transplant of *P. lanceolata* seedlings to individual microcosms, small (approximately 1.25 cm^3^) sections of ½GB5 agar containing abundant MFRE hyphae and spores were placed adjacent to emerging roots. Experimental microcosms were sealed with Parafilm and the ‘belowground’ agar portion of each plate was wrapped in aluminium foil to reduce light penetration into agar media. These microcosms were maintained in 16:8hr day:night conditions at a constant temperature of 25°C.

### Experimental microcosms

Using monoxenic *P. lanceolata*-MFRE microcosms (below), we established three experiments to test our hypotheses:

i. **To investigate the capability of MFRE to assimilate and transfer N to the host plant from diverse sources** we conducted an ‘*N source experiment’* whereby different ^15^N-labelled isotopes of compounds abundant in soils and transferred to plants by AM fungi (Hawkins *et al*., 2000; Reay *et al*., 2019) (ammonium, nitrate, urea, glycine; 1 mg.ml^-1^) were supplied to MFRE mycelium and subsequently measured in plant tissues while plant-derived C was traced into MFRE mycelium (Fig 1a).
ii. **To assess MFRE ^15^N source preference**, we conducted a ‘*fungal choice experiment’* whereby microcosms were simultaneously labelled with all four sources of N as the ‘N source experiment’, providing a choice of N source to MFRE mycelium (1 mg.ml^-1^ per source; Fig 1b).
iii. **To determine the effect of substrate N concentration variability** on MFRE-plant nutrient exchange and the fate of organic C bound within complex organic N sources, three different nutrient media treatments were employed in an ‘*N concentration experiment’* (Fig 1c). Each treatment was based on ½GB5 but with inclusion of differing quantities of N, concentrations being relevant to previous experimental systems (total N in media of experiments i and ii is equivalent to the ‘High N’ treatment of experiment iii) as well as a limestone grassland in the Peak district (Horswill *et al*., 2008; ‘Low N’). Treatments comprised: ‘High N’ (187.4 µg.g^-1^N), ‘Medium N’ (93.7 µg.g^-1^N), and ‘Low N’ (25 µg.g^-1^N) (Full nutrient composition in table S3).

**Figure 1.**
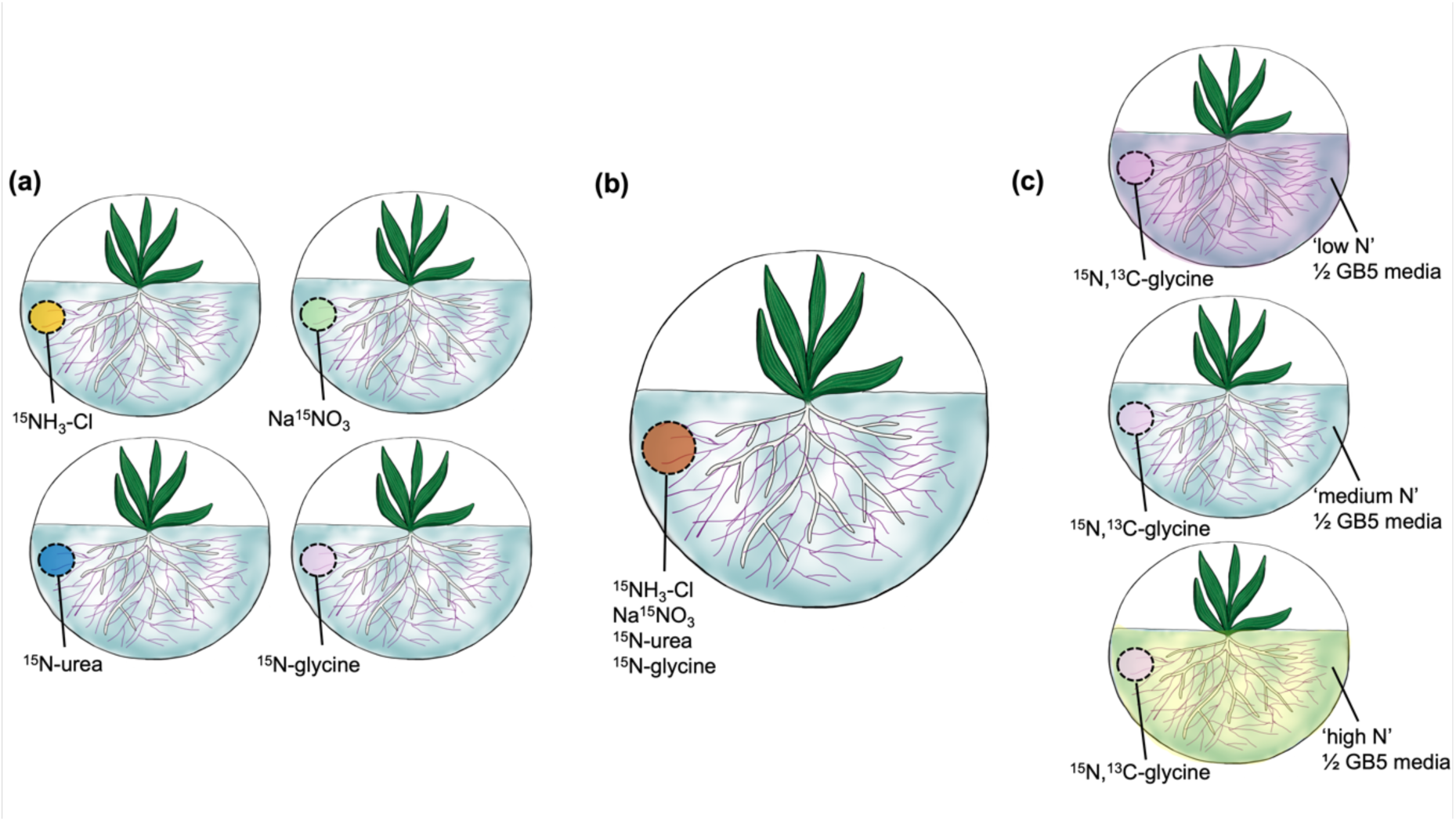
Monoxenic microcosm experiments. For tracing movement of C and N between the Mucoromycotina fine root endophyte fungal isolate *Lyc-1* and *Plantago lanceolata,* **(a)** *‘N source’* experiments where well contains one of ^15^NH_3_Cl, Na^15^NO_3_, ^15^N-urea or ^15^N-glycine; **(b)** *‘Fungal choice*’ experiments where well contains an equal mixture of all sources in (a); **(c)** *‘N concentration’* experiments where all microcosms are supplied with ^15^N/^13^C-glycine with media treatments comprising ‘high’ (187.4 µg g^-1^), ‘medium’ (93.7 µg g^-1^), and ‘low’ (25 µg g^-1^) N content of ½ GB5 agar (Full composition in table S3). Headspace of microcosms in (a) and (c) were additionally supplied with a 0.25 MBq pulse of ^14^CO_2_ to track movement of plant photosynthates to MFRE mycelium.

In all experiments, monoxenic microcosms (Fig. 1) were established using *Plantago lanceolata* seedlings (Yellow Flag Wildflowers, Gloucester, UK) with MFRE (*Lyc-1)* mycelium introduced from axenically-grown stocks (see above). *P. lanceolata* is a non-leguminous, mycorrhizal forb, common across a range of diverse habitats (Medina-van Berkum *et al*., 2024) and with a wide distribution (Penczykowski *et al*., 2021). It is a commonly used plant model for AM studies (Pankoke *et al*., 2015) with the relatively small size, propensity for mycorrhization (identified as hosting MFRE in wild-collected plants; Walker *et al*., 2018) and rapid development making it a very tractable and ecologically relevant plant species for our experiments. 140 mm sterile triple vented Petri-dishes were filled with ∼60 mL of ½GB5, or one of the three treatments in N concentration experiments, poured on a gradient that allowed for plant development in an upright position. *P. lanceolata* seedlings sterilised in a 4.5% sodium hypochlorite solution were germinated on flat ‘nursery’ plates of ½GB5 under 16:8hr (day:night) at room temperature. Seven days after sterilisation, individual germinated seedlings were transferred to experimental microcosms under sterile conditions.

### Colonisation of P. lanceolata roots by MFRE and mycelial growth

After growing plants and fungi together in microcosms for seven weeks, ‘*N concentration’* experiment plants were harvested and approximately 25% of the root system from each microcosm were stained using methods modified from Vierheilig *et al*., 1998. Quantification of colonisation by MFRE was not possible for the ‘*N source*’ and ‘*fungal choice*’ experiments due to the limited root biomass. Briefly, roots were placed into a 10% solution of KOH for 1hr at 70°C, rinsed in tap water, placed in ink-vinegar stain (5% Pelikan Brilliant Black, 5% acetic acid, 90% d.H_2_O) for 1hr, then rinsed again and placed in 1% acetic acid overnight to de-stain. The stained root material was mounted on slides in PVLG (Polyvinyl-Lacto-Glycerol) and colonisation counted under 40 x objective magnification (Ceti Max II; Medline Scientific, Chalgrove, UK). Representative images (Fig. 4) were obtained under 100 x objective magnification (Leica DM6; Leica Microsystems, Wetzlar, Germany).

We measured the two-dimensional area of MFRE extraradical mycelium in each microcosm of the ‘N concentration’ experiment weekly from the point of inoculation until the systems were harvested seven weeks later. The outline of the furthest extent of MFRE mycelial networks (Fig. 5d) were digitized and mycelial area determined using ImageJ (v1.53a; Schneider *et al*., 2012).

### ^15N^, ^13^C, and ^14^C isotope tracing

1. ‘*N source’ experiment*: Seven weeks after seedlings were placed in individual microcosms, a ∼2.5ml well was dug into the agar near to the margins of the MFRE mycelium, away from plant roots, filled with 100 µL of a 1 mg mL^-1^ solution of a single ^15^N-labelled compound (total 0.1 mg ^15^N labelled compound per plate; one of ammonium chloride (^15^NH_4_Cl, ≥ 98% atom % ^15^N, 27.53 µg ^15^N; Sigma-Aldrich), sodium nitrate (Na^15^NO_3_, ≥ 98% atom % ^15^N, 17.44 µg ^15^N; Sigma-Aldrich), glycine (C₂H₅^15^NO₂, ≥ 98% atom % ^15^N,19.72 µg ^15^N; Sigma-Aldrich) and Urea (CH₄^15^N₂O, ≥ 98% atom % ^15^N, 48.36 µg ^15^N; Sigma-Aldrich) and solidified with ½GB5 media. To control for diffusion of the ^15^N solution into the agar and subsequent direct plant assimilation, non-fungal control microcosms were also established. (n = 10 control microcosms for each ^15^N treatment apart from ^15^N-ammonium chloride which n = 9 controls due to microbial contamination). In total n = 20 (Na^15^NO_3_, ^15^N_2_-urea), n = 19 (^15^NH_4_Cl, ^15^N-Glycine).
2. ‘*Fungal choice’ experiment*: Seven weeks after inoculation with MFRE, wells were filled with 25 µL of a 4 mg.mL^-1^ solution of each N source used previously (i.e. ammonium chloride, sodium nitrate, glycine, and urea). These were applied in four treatments, with only one of the sources in each containing the ^15^N label. As such, each treatment comprised three unlabelled N sources and one ^15^N-labelled N source (0.1 mg compound per source, 0.4 mg compound in total per microcosm). Each well was backfilled with ½GB5 as previously described. The fungal ‘choice’ experiment comprised 10 fungal experimental microcosms. To control for diffusion of isotope through the agar medium, uninoculated control microcosms were established (n = 10 uninoculated microcosms per treatment). In total there were 20 microcosms established per treatment
3. *N concentration experiment*: Building on the observations from experiments i) and ii), seven weeks post inoculation, all microcosms of each N concentration treatment (‘High N’: 187.4 µg.g^-1^N, ‘Medium N’: 93.7 µg.g^-1^N, and ‘Low N’: 25 µg.g^-1^N) were labelled with a solution of 100 µl 1 mg.ml^-1^ ^15^N-glycine tracer added to wells cut into the agar portion of the microcosms and backfilled with ½GB5 media, as described above (n = 5 per treatment). To determine the fate of glycine bound-C, we labelled using stable ^15^N- (19.72 µg ^15^N) and to determine the fate of glycine bound-C in the presence of MFRE, we used ^13^C-labelled glycine (17.09 µg ^13^C per plate). Shoot ^15^N/^13^C concentrations were determined using IRMS. To control for isotope diffusion and non-MFRE mediated N/C distribution, non-fungal control microcosms were established (n = 5 uninoculated microcosms per treatment).

In ‘N source’ and ‘N concentration’ experiments, immediately after ^15^N/^13^C addition into wells, the surface of the agar portion of the microcosm was covered with a clear PVC sheet and sealed with anhydrous lanolin. A 0.25 MBq ^14^CO_2_ pulse was liberated into the headspace of sealed plates from 6.75 µl ^14^C-labelled sodium bicarbonate (2.14 GBq/mmol) by the addition of 2 ml 90% lactic acid. Microcosms were incubated for 24 hrs to allow for ^14^CO_2_ fixation and movement of ^15^N (and ^13^C) and ^14^C between plants and MFRE. At the end of the labelling period, 2 ml 2M KOH was introduced into small containers within the microcosms to absorb any remaining ^14^CO_2_. After 1 hr, all plant materials were removed carefully from the agar, separating plant shoots from roots, and removing as much excess agar from root material as possible prior to freeze-drying. To assess ^14^C transfer to MFRE the agar (containing MFRE fungal mycelium in all microcosms apart from uncolonized controls) was also freeze-dried and homogenised. 10-30 mg freeze-dried agar was weighed into CombustoCones (Perkin Elmer, Beaconsfield, UK) prior to sample oxidation (Sample Oxidiser 307, Perkin Elmer, Beaconsfield, UK) and ^14^C quantification via liquid scintillation counting (Packard Tri-Carb 4910TR, Perkin Elmer, Beaconsfield, UK). Total carbon (^12^C + ^14^C) fixed by the plant and transferred to MFRE within the agar was calculated as a function of the total volume and CO_2_ content of the labelling chamber and the proportion of the supplied ^14^CO_2_ label fixed by the plants. The difference in carbon between fungal and non-fungal plants is equivalent to the total C transferred from plant to MFRE within the fungal microcosms, assuming no alteration in plant root C exudation under fungal colonisation. Total carbon assimilated by the plant was calculated using the following equations modified from Hoysted *et al*. (2023):

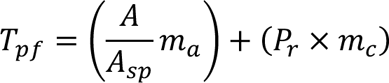

Where *T_pf_* = Transfer of carbon from plant to fungus, *A* = radioactivity of the agar tissue sample (Bq); *A_sp_* = specific activity of the source (Bq Mol^−1^), *m_a_ =* atomic mass of ^14^C, *P_r_* = proportion of the total ^14^C label supplied present in the agar tissue; *m_c_* = mass of C in the CO_2_ present in the labelling chamber (g) (from the ideal gas law):

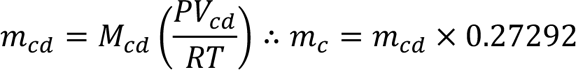

where *m_cd_* is mass of CO_2_ (g), *M_cd_* is molecular mass of CO_2_ (44.01 g.mol^-1^), P is total pressure (kPa); *V_cd_* is the volume of CO_2_ in the chamber (0.000049m^3^); *R* is the universal gas constant (J.K^-1^.mol^-1^); *T*, absolute temperature (K); *m_c_*, mass of C in the CO_2_ present in the labelling chamber (g), where 0.27292 is the proportion of C in CO_2_ on a mass fraction basis. To determine the amount of C transfer to agar that was mediated by MFRE alone, the average concentration of ^14^C in non-fungal controls was subtracted from the ^14^C concentration in individual experimental microcosms.

In all experiments, all plant shoots were harvested and freeze-dried 24 hr after isotope addition. Between 0.1 and 5mg freeze-dried shoot tissue were measured into tin capsules (Sercon, Crewe, UK) and the abundance of ^15^N in samples determined by IRMS (Isotope Ratio Mass Spectrometry) using an ANCA GSL 20-20 Mass Spectrometer (Sercon PDZ Europa 2020 Isotope Ratio Mass Spectrometer coupled to a PDZ ANCA GSL preparation unit). Data were collected as atom %^15^N and as %N using un-labelled control plants for background detection. Plant tissue concentration of ^15^N was calculated using the following equations from Hoysted *et al*. (2023):

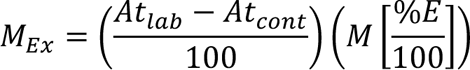

Where *M_Ex_* is mass (excess) of ^15^N in samples (g), *At_lab_* is atom percentage of ^15^N in the experimental microcosms, *At_cont_* is the atom percentage of ^15^N in unlabelled control plant material, this was generated by growing *P. lanceolata* seedlings in microcosms as described above but with no isotope labels added into the systems. *M* is the sample biomass (g) and *%E* is the total percentage of N. This was then converted to µg to obtain concentration per mg of plant tissue and then further expressed per g of plant biomass ([^15^N]). The average [^15^N] of non-fungal control microcosms for each ^15^N treatment was then subtracted from the [^15^N] for each experimental microcosm within that treatment.

### Statistical Analyses

All statistical analyses were conducted using (R Development Core Team, 2023) and R (v2023.3.0.386, R Core team, 2023), using packages ‘dplyr’ (v1.1.2, Wickham *et al*., 2023), ‘car’ (v3.1-2, Fox and Weisberg, 2019), ‘rosetta’ (v0.3.12, Peters and Verboon, 2023), ‘stats’ (v4.3.0, R Core team, 2023), ‘agricolae’ (v1.3-5, Mendiburu and Yaseen, 2020)] Isotope tracing data were analysed using analysis of variance (ANOVA) with post hoc Tukey testing (as indicated). Data were checked for normality and homogeneity of variance. Where assumptions were not met, either a square root or logarithmic transformation was performed (Table S1), or a non-parametric test Kruskal-Wallis with Dunn’s post hoc test (as indicated) was conducted. Plant biomass was compared between Fungal and non-Fungal plants using either a student’s T-test, or Wilcoxon signed-rank test where assumptions of normality and homogeneity of variance were not met. Hypha area growth data were analysed using a two-way repeated measures ANOVA with a bonferroni correction. Figures were created in R (v2023.3.0.386, R Core team, 2023) using the ‘ggplot2’ package (v3.4.2, Wickham, 2016).

## Results

### i) ‘N source’ experiment (**Fig. 1a**)

#### Plant growth

There was no effect of MFRE colonisation or ^15^N source type on root biomass (Fig. S1a; Table S1), although shoot biomass (Fig. S1b) was overall greater for MFRE inoculated plants compared to asymbiotic counterparts (ANOVA: *F_1,110_* = 7.05, *p* < 0.01), however no differences were observed in biomass between inoculated and uninoculated plants of the same treatment (Tukey’s HSD: *p* > 0.05; Fig. S1b). The type of ^15^N source supplied had some effect on shoot biomass (ANOVA: *F_3,110_* = 3.29, *p* < 0.05). There was no interaction between the factors (Table S1).

#### MFRE-mediated plant ^15^N concentration

In ‘*N source*’ experimental microcosms, the type of ^15^N source available to MFRE influenced ^15^N concentration in plant shoots (ANOVA: F*_3,74_*=3.7022, *p* < 0.05; Fig. 2a). Plants in microcosms where MFRE was supplied with ^15^N-glycine accumulated more ^15^N in the shoots than microcosms treated with either ^15^N-sodium nitrate or ^15^N_2_-urea (Tukey’s HSD: *p* < 0.05) but not those supplied with ^15^N-ammonium chloride.

**Figure 2.**
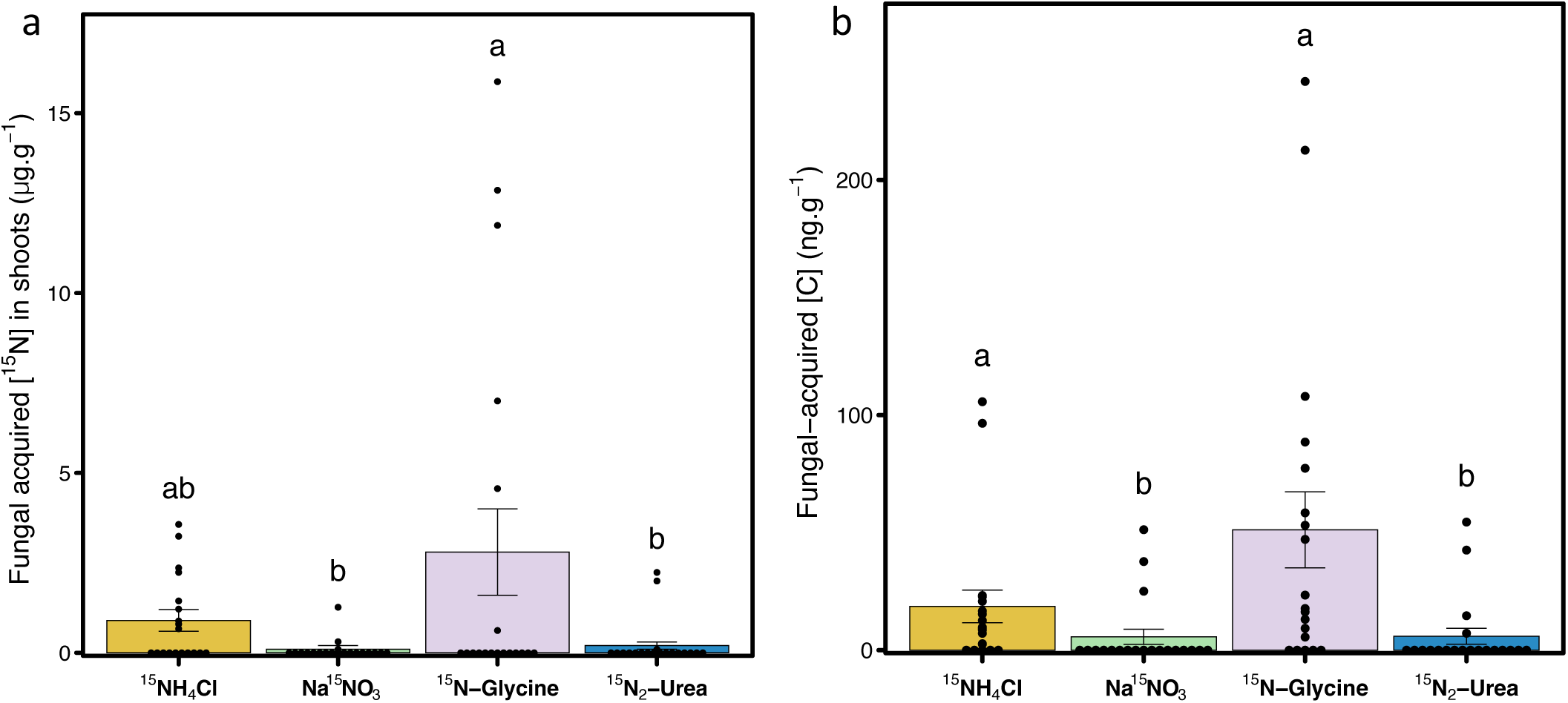
Nutrient tracing data from ‘N source’ experiment. a) Mean fungal-acquired ^15^N concentration of shoots. Different letters denote significant difference (Tukey’s HSD: p<0.05). n = 20 (Na^15^NO_3_, ^15^N_2_-urea), n = 19 (^15^NH_4_Cl, ^15^N-Glycine). Error bars indicate ±SE. **b)** Plant-derived C concentration in MFRE hyphae. n = 20 (Na^15^NO_3_, ^15^N_2_-urea), n = 19 (^15^NH_4_Cl, ^15^N-Glycine). Different letters denote significant difference (Dunn’s post hoc: p<0.05). Error bars indicate ±SE.

#### Plant-to-fungus C transfer

Quantification of fungal-acquired carbon in the ‘*N source*’ experiment was determined by calculating the mean ^14^C concentration of media in non-fungal microcosms (representative of diffusion and root exudation) and subtracting that value from the ^14^C concentration in individual fungal microcosms. ^12^C was determined using established equations (see methods) and added to ^14^C to provide ‘total C’. The trends we observed in plant-to-MFRE C transfer (Fig. 2b) mirror – and are stronger than – those observed for MFRE-mediated ^15^N transfer to plants (Fig. 2a). ^15^N source is a significant driver of the amount of plant C transferred to MFRE (Kruskal-Wallis: d.f. = 3, X^2^ = 20.256, *p* < 0.001), with microcosms treated with ^15^N-glycine or ^15^NH_4_Cl transferring significantly more C to MFRE than microcosms treated with either ^15^N_2_-urea or Na^15^NO_3_.

### ii) ‘Fungal choice’ experiment (Fig. 1b)

#### MFRE-mediated plant ^15^N transfer and assimilation

When a choice of N source was supplied to MFRE, ^15^N concentration in host plant shoots was strongly influenced by which compound contained the ^15^N label (ANOVA: *F_3,35_* = 4.3933, *p* < 0.01; Fig. 3). Plants in microcosms where the glycine or ammonium chloride components of the N source mixture supplied to MFRE were labelled with ^15^N accumulated more ^15^N in shoots than microcosms where the labelled ^15^N source was Na^15^NO_3_ (Tukey’s HSD: *p* < 0.05). Those microcosms supplied with urea as the ^15^N labelled components had intermediate shoot [^15^N] (Fig. 3).

**Figure 3.**
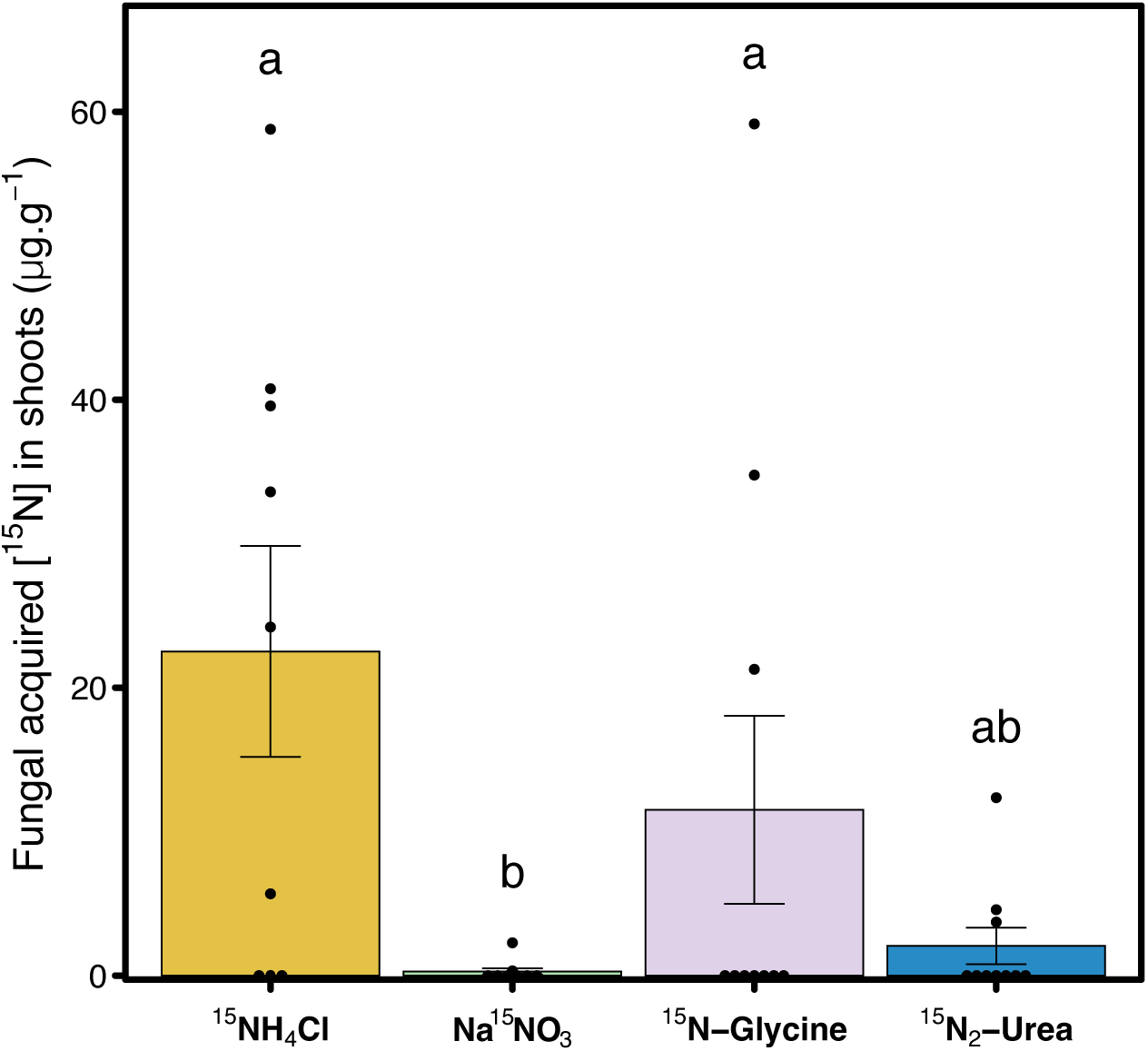
Fungal-acquired ^15^N content of shoots in ‘fungal choice’ experiment. Data presented as concentration. Different letters denote significant difference (Tukey’s HSD: p<0.05). **b**) n = 10 per treatment apart from ^15^NH_4_ which n = 9. Error bars indicate ±SE.

### iii) ‘N concentration’ experiment (Fig. 1c)

#### Colonisation by MFRE and extraradical MFRE hyphal growth

To study the impact of N-concentration on MFRE colonisation we quantified the colonisation of *P. lanceolata* roots by MFRE in each microcosm after 7 weeks of growth in monoaxenic systems with different levels of available ammonium and nitrate N (Table S3). Roots inoculated with MFRE were colonised by abundant fine (<1.5 µm diameter) hyphae which displayed typical MFRE morphology. We observed irregular branching (Fig. 4a-f), small (1-2 µm diameter) swellings (Fig. 4b-d), larger vesicle-like swellings (∼5 µm diameter; Fig. 4b), and smaller swellings at hyphae termini (∼6.54 µm diameter; Fig. 4f) within root cells, as previously observed in monoxenically grown plants Hoysted *et al*., (2019 and 2023). % Root colonisation by MFRE in the Low N treatment was significantly lower than in the Medium N or High N treatments (Fig 5a, ANOVA: *F_2,39_* = 6.5694, p < 0.01; Tukey’s HSD: p < 0.05). The presence of vesicle-like hyphal swellings was also lower in the Low N treatment than in the Medium or High N treatments (Fig. 5b, ANOVA: *F_2,39_* = 10.561, p < 0.001). The surface area of extraradical MFRE mycelium extending beyond the root (Fig. 5c) was greater at all time points measured in Low- and Medium N treatments compared to High N (ANOVA: *F_3.81,74.37_* = 14.856, p < 0.001).

**Figure 4.**
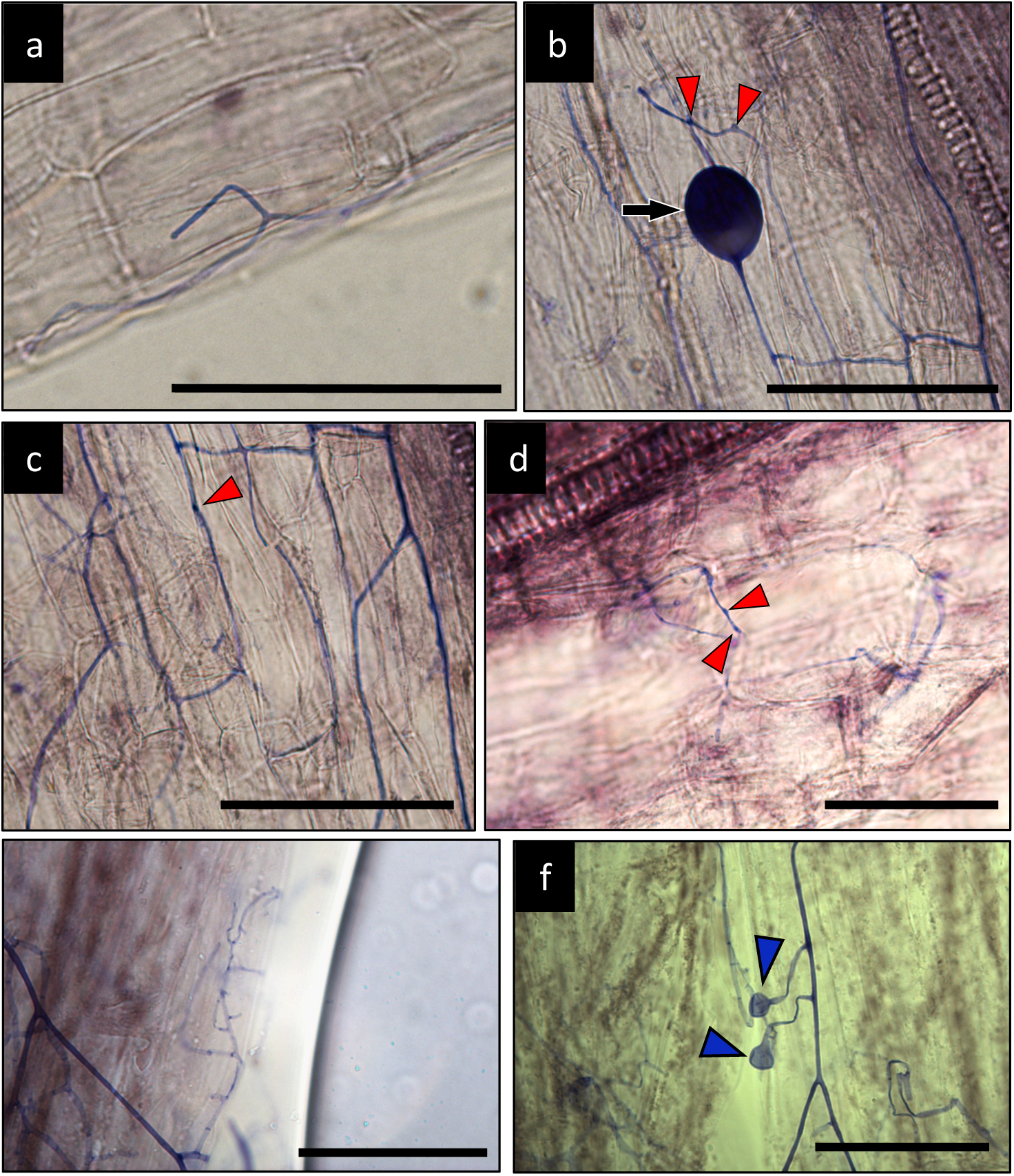
Ink-stained roots of *P. lanceolata* colonised by MFRE (in purple) showing fine (<1.5µm diameter) branching hyphae within cells. (a-e) including those with small swellings (b-d; red wedges) as well as larger vesicles (b; black arrow) and terminal swellings (f; blue wedges). All images are from samples taken from monoxenic microcosms 7 weeks post inoculation with MFRE. Scale bars = 50µm.

#### MFRE-to-plant ^15^N and plant-to-MFRE C transfer

Plants growing in Low N microcosms (Full composition in table S3) contained significantly greater MFRE-acquired [^15^N] in shoots than those grown in Medium- or High N treatments (Kruskal-Wallis: X^2^= 9.9668, d.f. = 2, p < 0.01; Dunn test: p < 0.05) (Fig. 6a). There were no significant differences in plant-fixed [C] (ng.g^-1^) transferred to MFRE mycelium, regardless of N concentration in the media (Fig. 6b; Table S1). Plant shoot ^13^C concentration (Figure 6c) was greater in asymbiotic plants in Medium- and Low-N treatments compared to MFRE-colonised plants within the same treatment (ANOVA: *F_1,50_* = 20.6347, p < 0.001). Although asymbiotic plants were not affected by the N content of the media, MFRE-colonised plants accumulated more ^13^C in shoot tissues in the High N media compared to the Low N media while an intermediate amount of ^13^C accumulated in shoots of plants grown in Medium N media (ANOVA: *F_2,50_* = 5.0497, p <0.01; Fig 6c).

## Discussion

Our experiments show direct C-for-N exchange between MFRE (*Lyc-*1 isolate) and the AM host plant *Plantago lanceolata*(Stewart, 1996; Pankoke *et al*., 2015; Pel *et al*., 2018) *in vitro*. Although not all microcosms in our experiments demonstrated C-for-N transfer between MFRE and host plants (likely due to the relatively short time frame of our isotope tracing period compared with other tracer studies, e.g. Hawkins *et al*., 2000; Govindarajulu *et al*., 2005; Thirkell *et al*., 2020), there was a preferential transfer and assimilation of ^15^N derived from glycine and ammonium than from nitrate or urea (Fig. 2a, 3a) into host plant tissues via MFRE symbionts. This corresponded to enhanced photosynthate allocation to the fungus (Fig. 2b).

To date, demonstration of resource exchange between MFRE and host plants in the absence of other soil microbes has been limited to a handful of plant species. The only other study of an angiosperm being that of Hoysted *et al*. (2023) where MFRE transferred N to the legume *Trifolium repens* in return for plant-fixed carbon resources in the absence of other soil microbes. However, in the *Trifolim* study the only source of ^15^N tracer was ^15^NH_4_Cl. Further, as a legume, it is likely that *T. repens* is less reliant on fungal symbionts for N acquisition than non-leguminous plants as N is typically supplied to the plant by N-fixing bacterial symbionts. Our data expand the known range of sources from which MFRE can access and transfer N to a non-leguminous plant host. Given the range and preferences demonstrated by MFRE in our experiments, the possibility of functional complementarity with AM fungi in dual colonisations whereby AM fungi play a primary role in supplying host plants with P (Johansen *et al*., 1996; Toussaint *et al*., 2004) while MFRE supply N from a variety of sources (Field *et al*., 2019; Hoysted *et al*., 2019, 2023), remains open.

The greater MFRE-mediated assimilation and transfer of glycine- and ammonium-^15^N to *P. lanceolata* compared to the other ^15^N sources supplied occurred regardless of whether the ^15^N-ammonium or ^15^N-glycine tracer was the only ^15^N source provided to the extraradical MFRE mycelium or whether these sources were part of a mixture (Fig. 5b). The preference for glycine-^15^N by MFRE in our experimental systems contrasts with the tendency of many AM fungal species to assimilate N from inorganic sources for transfer to host plants (Johansen *et al*., 1996; Toussaint *et al*., 2004). When presented with ^15^N-glycine in non-sterile microcosms, four species of AM fungi showed no direct transfer of the tracer to host plants (Hodge, 2001). As obligate biotrophs, AM fungi lack the molecular toolkit required to degrade organic molecules, such as the extracellular proteases that are commonly found in saprotrophic ericoid and ectomycorrhizal fungi (Jin *et al*., 2012). In contrast, because MFRE can be isolated from host plants and cultured in a free-living state in axenic conditions (Field *et al*., 2015; Hoysted *et al*., 2023), MFRE must possess at least some degradative capabilities to maintain mycelial growth in the absence of a host plant. Our key findings that MFRE preferentially assimilate and transfer glycine-derived N to the host plant in return for proportional photosynthetic C, and that MFRE-associated plants are inhibited in their acquisition of ^13^C tracer from glycine while transfer of glycine-^15^N is maintained, suggest that MFRE retain and metabolise glycine-derived C skeletons, liberating ammonium-N for transfer to host plants. This provides strong evidence of independent exogenous C acquisition capabilities of MFRE, even when associated with a living host plant. The acquisition of N from some small organic compounds by AMF has been demonstrated (Matsumura *et al*., 2013) but to date there is no convincing evidence of AM fungi utilising C from organic sources (Hodge and Storer, 2015).

**Fig 5.**
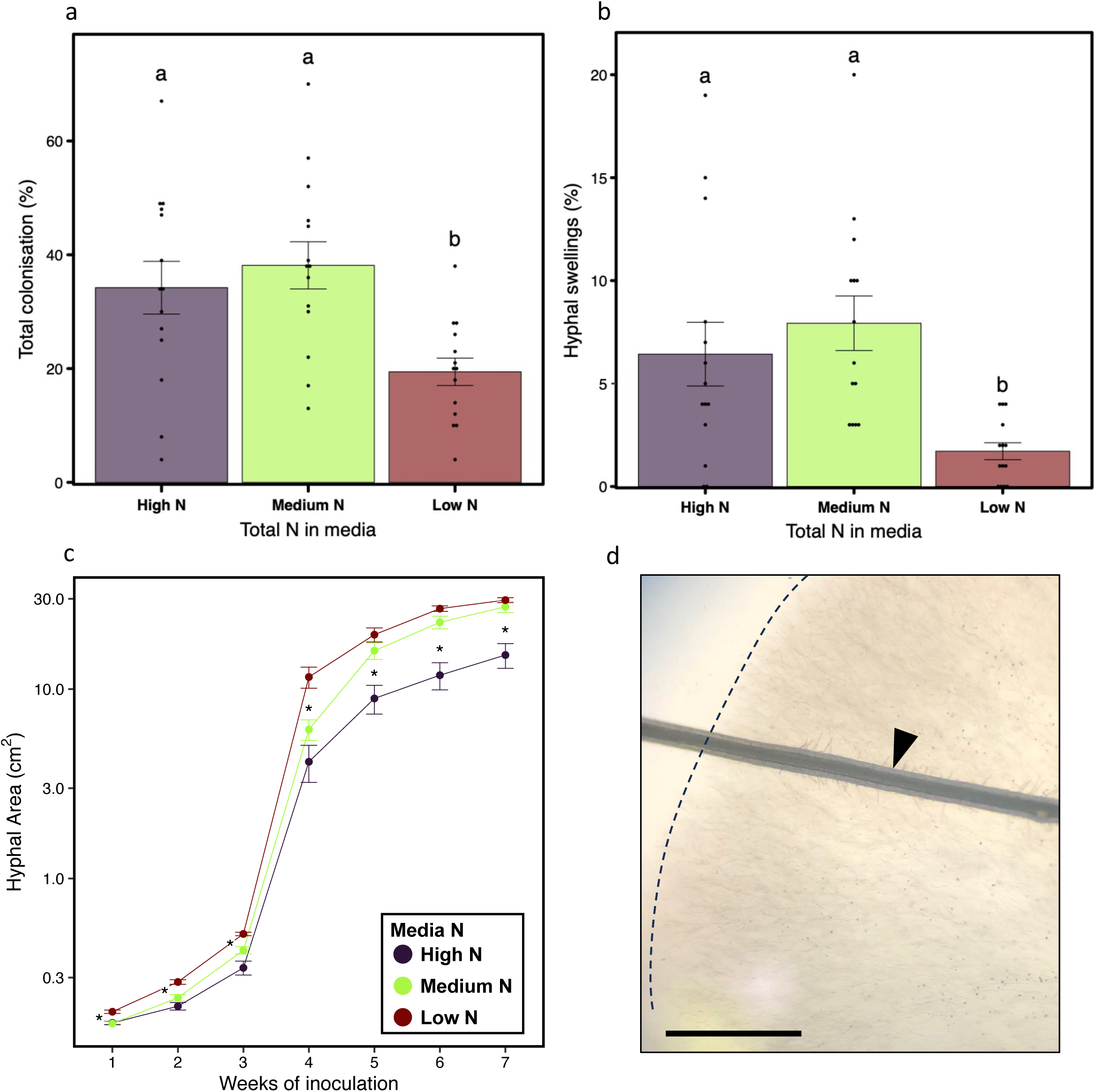
Colonisation of *Plantago* roots by MFRE structures (a & b), and hyphal area beyond roots (c). **a)** Total colonisation by fungal structures. n=14 per N treatment. **b)** Colonisation by spherical ‘vesicular’ structures n=14 per N treatment. **c)** Growth of extraradical MFRE mycelium in the ‘N concentration’ experiment. n=14 per N treatment per time point. **b**) Image of *P. lanceolata* root on ½GB5 medium (black wedge) and dense area of MFRE hyphae with defined growing edge (dashed line). Scale bar = 500µm.

There was little-to-no transfer of ^15^N derived from nitrate to host plants by MFRE in any of our experiments, regardless of whether nitrate was included as part of a mixture of N sources or was the only source of N added to the microcosm (Fig. 5a and b). This contrasts with AM fungi which frequently assimilate nitrate-N and transfer it to host plants (Toussaint *et al*., 2004; Calabrese *et al*., 2016) and possess nitrate reductases within their genomes (Kaldorf *et al*., 1994). Compared to the other N sources in this study, the relative bioavailability of nitrate-N to both plants (Noguero and Lacombe, 2016) and AM fungi (Bago *et al*., 1996; Tian *et al*., 2010) within the rhizosphere may explain the lack of MFRE-mediated nitrate-N transfer to host plants as assimilation of nitrate-N would represent a relatively more competitive niche for MFRE to exploit. As such, a plausible hypothesis would be that MFRE do not possess the requisite molecular or metabolic capacity to exploit soil nitrate pools having instead evolved capabilities to acquire and metabolise both N and C from organic sources. While also a small inorganic molecule, ammonium-N – in contrast with nitrate-N – was transferred to plants in amounts comparable to glycine-N. This may be evidence that MFRE assimilates N from soils as ammonium in a similar manner to AM fungi (Marzluf, 1997) however more research is necessary to confirm this hypothesis.

Although MFRE transferred ^15^N from glycine to host plants, the amount of ^15^N delivered to host plants from urea, the other organic N source available to the fungi in our experiments, was much lower. This is probably due to the chemical nature of urea; urea is a stable molecule, with a half-life of over three years in solution (Amtul *et al*., 2002) only after which time it degrades to produce ammonium ions. Urease enzymes which catalyse this reaction are produced by a range of soil microbes (Rana *et al*., 2021) including AM fungi (Jin *et al*., 2012). It may be that MFRE do not produce these enzymes to utilise this substrate, instead scavenging ammonium-N – which our results suggest is readily utilised by MFRE (Fig. 5) – from decomposition of urea by other soil-borne microorganisms. The addition of urease-producing soil bacteria to MFRE-only experimental systems, or the development of soil-based systems replete with a rhizosphere microbiome to investigate MFRE-plant symbioses is now needed to determine whether this is the case.

In AM symbioses, plant reliance on fungal-acquired nutrients is highly context dependent (Kiers *et al*., 2011). In particular, the availability of nutrients within the environment can play a large role in determining the stoichiometry of P and N transfers from AM fungi to host plants and shifts in fungal community composition (Kiers *et al*., 2011; Jach-Smith and Jackson, 2018; Lilleskov *et al*., 2019; Lekberg *et al*., 2021). To test whether the nutritional role of MFRE is similarly plastic, we investigated the N transfer from MFRE-to-plant across a range of N concentrations supplied to the growth media. Given the preference for assimilation and transfer of N from glycine observed in our other experiments, we supplied the MFRE mycelium with ^15^N/^13^C labelled glycine in modified ½GB5 media (Table S3). We found that host plants relied on MFRE for a greater proportion of their ^15^N assimilation when grown on reduced N availability media (Fig. 6a). The low N media stimulated growth of a larger extraradical MFRE mycelial network (Fig. 5c) and reduced presence of structures typical of colonisation within host roots (Fig. 5a and b) than the other media treatments, suggestive of an explorative, foraging growth strategy being deployed by the fungus. In contrast, there was no increase in C allocation from plant hosts to MFRE under the same limited N growth conditions (Fig. 6b). However, there was a corresponding decrease in plant acquisition of ^13^C from glycine in low N conditions (Fig. 6c), indicating assimilation and sequestration of ^13^C (and ^12^C) from glycine by MFRE. C acquisition by MFRE from an exogenous source could offset ‘costs’ associated with increased transfer of N by MFRE to hosts (Kiers *et al*., 2011) in low-N environments. The increased area of MFRE hyphae beyond plant roots in low N media may explain the greater transfer of ^15^N observed under the same conditions as extension of foraging fungal hyphae would increase the likelihood of their encountering the isotope tracer, increasing the amount of tracer transferred to hosts despite no greater root colonisation being recorded.

**Figure 6.**
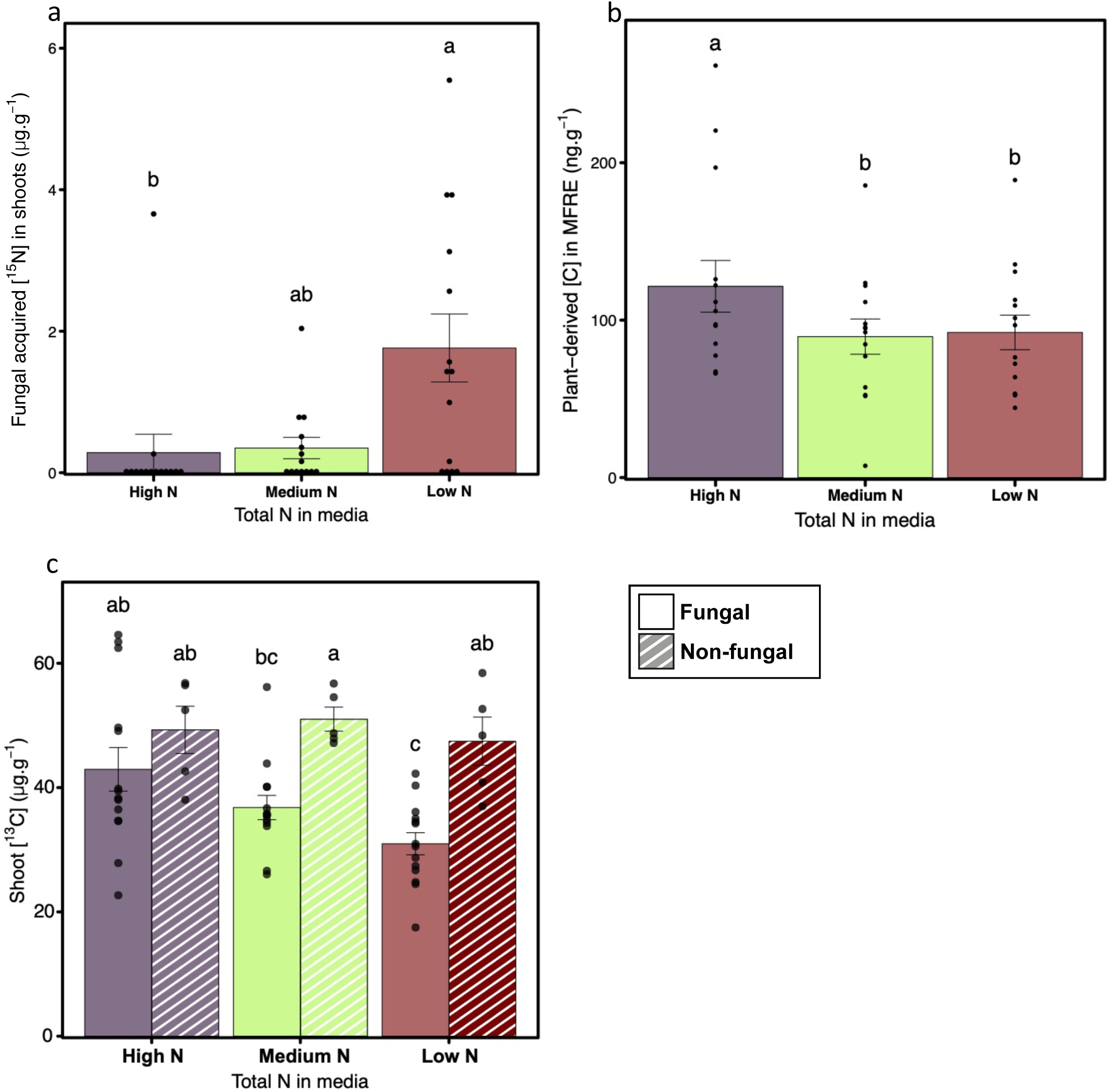
Nutrient tracing data for the ‘N concentration’ experiment. **a)** concentration of fungal-derived ^15^N from labelled glycine in plant shoots. Different letters denote significant difference (Dunn’s post hoc: p<0.05). n = 14 per treatment. **b**) Plant-derived C allocation to agar compared between fungal and non-fungal microcosms. Different letters denote significant difference (Tukey’s HSD: p<0.05). n = 9 per fungal treatment, n = 5 per non-fungal treatment. **c**) Plant glycine-derived ^13^C concentration compared between fungal (solid bars) and non-fungal (hatched bars) microcosms. Different letters denote significant difference (Tukey’s HSD: p<0.05). n = 9 per fungal treatment, n = 5 per non-fungal treatment. Error bars indicate ±SE.

Our finding that MFRE preferentially transfer N from glycine host plants (Fig. 2a and 3) contrasts with our hypothesis that inorganic N would be preferred owing to its relatively simple structure, resulting in more energy efficient assimilation and metabolism compared to more complex organic compounds. However, our finding that MFRE reduce the assimilation of glycine-derived ^13^C by plants (Fig. 6c) while simultaneously enhancing assimilation of glycine-N (Fig. 6a) and receiving plant-derived C (Fig. 6b) is consistent with the capacity of MFRE to supplement its plant-fixed C nutrition with C uptake from environmental sources. Access to nutrients bound up in organic compounds potentially provides MFRE with a competitive edge in a crowded symbiotic marketplace, providing the fungi with an advantage when there is not a ready supply of plant-derived C. As such, MFRE may preferentially assimilate organic compounds in the soil as these provide not only N, an exchangeable commodity in plant-fungal symbioses, but also C which can be used by the fungus to supplement its supply of plant-fixed C. If this is the case, then assimilation of organic compounds could ultimately provide more benefit to MFRE than an inorganic N source. Our experiments support this hypothesis, showing that regardless of N availability in the growth media, MFRE obtained similar amounts of plant-fixed C (Fig. 6b) despite greater N being transferred to host plants from glycine. Such physiological plasticity is likely to facilitate persistence of MFRE alongside, and in competition with, other plant endophytic fungi for space and resources within host plant roots.

It is important to note that our experimental systems do not represent the full complexity of soil ecosystems. In nature, plant-MFRE symbioses appear to occur in most scenarios that have been investigated (Albornoz *et al*., 2021), encompassing many other abiotic factors including variable light conditions, nutrient availability and access to water. As such our findings should not be generalised to all plant-MFRE symbioses in all environmental scenarios; crucially the data presented excludes the potential for MFRE-microbial interactions, such as those between AM fungi and soil bacteria (Jiang *et al*., 2021). Such interactions may facilitate the assimilation of a broader range of N sources by MFRE. Nevertheless, our work here represents an important starting point for the exploration of the broader ecological and physiological significance of MFRE to develop a more holistic understanding of plant-fungal symbioses.

## Acknowledgements

Funding: This work was funded by an ERC consolidator grant (MYCOREV, project ref: 865225) to K.J.F., T.J.D. and S.P. and a NERC grant to K.J.F. (NE/S009663/1). E.D. is funded by a NERC ECORISC PhD studentship. We thank the De Laszlo Foundation for generously supporting PhD student research. Author contributions; N.H. and K.J.F. conceived and designed the experiments; N.H., E.D., A.W., and K.J.F performed the experiments and laboratory analyses; N.H. analysed and interpreted the data; N.H. wrote the manuscript with contributions from all other authors. All authors revised and approved the final manuscript.

