## Supplementaryinfo for "Preferential assimilation, metabolism, and transfer of organic nitrogen to host plants by Mucoromycotina ‘fine root endophytes’"

### Figure S1. Biomass

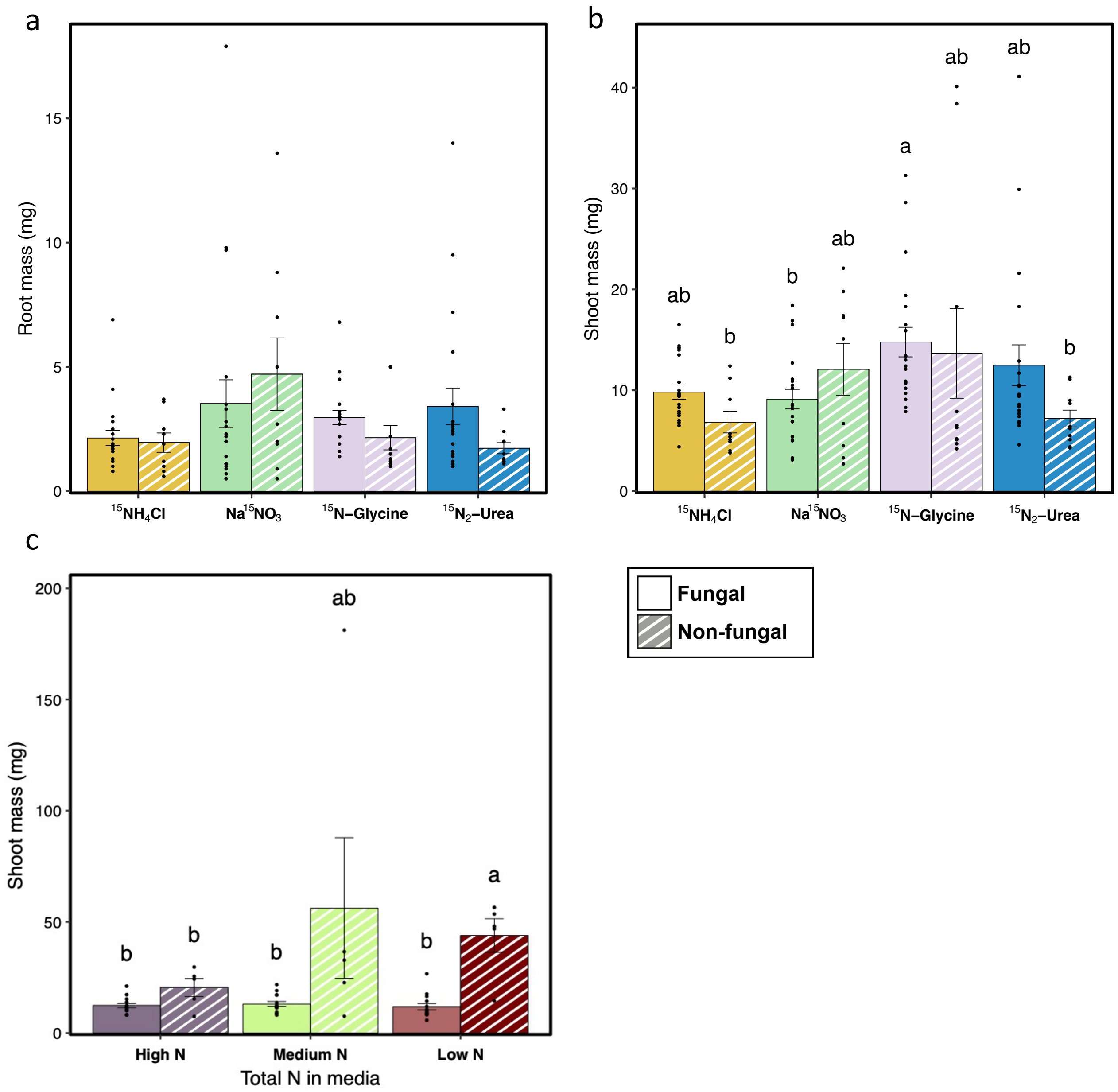


**Figure S1. Biomass (mg) of plant roots (a) and shoots (b & c)** in single 'N source' experiments **(a & b)**, and 'N concentration' experiments **(c)**. Shaded bars indicate microcosms not inoculated with MFRE and open bars indicate inoculation with MFRE. Error bars indicate ±SE, different letters denote significant difference (Tukey's HSD: $p<0.05$). **a** & **b**; n = 20 per fungal treatment, n = 10 per non-fungal treatment apart from $^{15}NH_4$ which n = 9) and **c)** n = 9 per fungal treatment, n = 5 per non-fungal treatment. Different letters denote significant difference (Tukey's HSD: $p<0.05$). Error bars indicate ±SE.

#### Figure S2. Total N

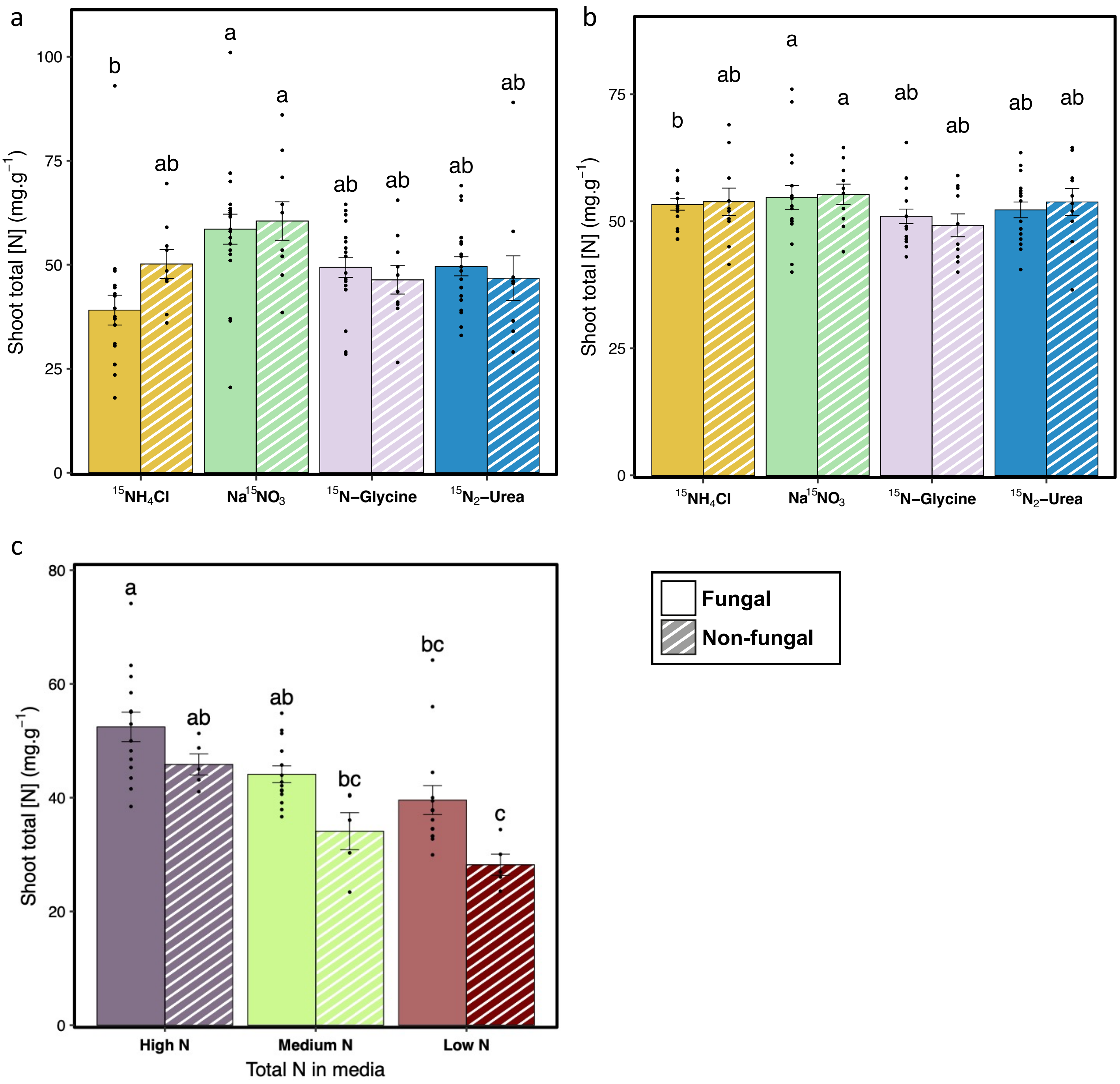


**Figure S2. Total N concentration of plant shoots** in ‘N source’ (**a**), ‘fungal choice’ (**b**), and ‘N concentration’ (**c**) experiments. **a)** n = 20 per fungal treatment, n = 10 per non-fungal treatment apart from $^{15}NH_4$ which n = 9). **b)** n = 17 per fungal treatment apart from $^{15}NH_4$ which n = 16, n = 10 per non-fungal treatment. **c)** n = 14 per treatment for fungal microcosms, n = 5 per treatment for non-fungal microcosm. Shaded bars indicate microcosms not inoculated with MFRE, open bars indicate inoculation with MFRE. Error bars indicate ±SE, different letters denote significant difference (Tukey's HSD: $p<0.05$).

#### Table S1. Statistical tests

##### SI Table 1 Statistical tests, results, and transformations performed on data

| Figure | Statistical test | Variable(s) tested | Transformation | P value | Post hoc test & p value |
|---|---|---|---|---|---|
| **3a** | ANOVA: d.f.=3,74, F=3.7022 | Shoot [$^{15}N$] (μg.g$^{-1}$) | Square root transformed | p < 0.05 | Tukey's HSD; p<0.05 |
| **3b** | Kruskal-Wallis; X$^2$=20.256, d.f.=3 | Plant-derived $^{14}C$ in MFRE containing media | N/A | p < 0.001 | Dunn's post-hoc: p<0.05 |
| **4** | ANOVA: d.f.=3,35, F= 4.3933 | Shoot [$^{15}N$] (μg.g$^{-1}$) | Square root transformed | p < 0.01 | Tukey's HSD; p<0.05 |
| **5a** | ANOVA; d.f.=2,39, F=6.5694 | Total Colonisation | N/A | p < 0.01 | Tukey's HSD: p<0.05 |
| **5b** | ANOVA; d.f.=2,39, F=10.561 | Vesicular Colonisation | N/A | p < 0.001 | Tukey's HSD: p<0.05 |
| **5c** | ANOVA; d.f.=2,39, F= 20.186<br>ANOVA; d.f.=1.91,74.37, F= 350.927<br>ANOVA; d.f.=3.81,74.37, F= 14.856 | N concentration<br>Time point<br>Interaction | N/A | p < 0.001<br>p < 0.001<br>p < 0.001 | Pairwise t-tests (Table S2) |
| **6a** | Kruskal-Wallis: d.f. = 2, X$^2$= 9.9668 | Shoot [$^{15}N$] (μg.g$^{-1}$) | N/A | p < 0.01 | Dunn's post-hoc: p<0.05 |
| **6b** | ANOVA: d.f = 2,39, F= 1.8301 | [C] in MFRE hyphae (ng.g$^{-1}$) | N/A | p > 0.05 | Tukey's post-hoc: p > 0.05 |
| **6c** | ANOVA; d.f.=1,50, F= 20.6347 | Fungal inoculation | N/A | p < 0.001 | Tukey's HSD: p<0.05 |
| | ANOVA; d.f.=2,50, F=5.0497 | N concentration | | p < 0.01 | |
| | ANOVA; d.f.=2,50, F=1.2897 | Interaction | | p > 0.05 | |
| **S1a** | ANOVA; d.f.=1,110, F=0.5905 | Fungal inoculation | N/A | p > 0.05 | N/A |
| | ANOVA; d.f.=3,110, F=2.2187 | $^{15}N$ treatment | | p > 0.05 | |
| | ANOVA; d.f.=3,110, F=1.2649 | Interaction | | p > 0.05 | |
| **S1b** | ANOVA; d.f.=1,110, F=7.0530 | Fungal inoculation | Log transformed | p < 0.01 | Tukey's HSD; p<0.05 |
| | ANOVA; d.f.=3,110, F=3.2880 | $^{15}N$ treatment | | p < 0.05 | |
| | ANOVA; d.f.=3,110, F=1.7085 | Interaction | | p > 0.05 | |
| **S1c** | ANOVA; d.f.=1,51, F=37.6797 | Fungal inoculation | Log transformation | p < 0.001 | Tukey's HSD: p<0.05 |
| | ANOVA; d.f.=2,51, F=0.7200 | N concentration | | p > 0.05 | |
| | ANOVA; d.f.=2,51, F=2.9633 | Interaction | | p > 0.05 | |
| **S2a** | ANOVA; d.f.=1,109, F=4.80 | Fungal inoculation | Log transformed | p < 0.05 | Tukey's HSD: p<0.05 |
| | ANOVA; d.f.=3,109, F=3.91 | $^{15}N$ treatment | | p < 0.01 | |
| | ANOVA; d.f.=3,109, F=1.52 | Interaction | | p > 0.05 | |
| **S2b** | ANOVA; d.f.=1,99, F= 7.26 | Fungal inoculation | Log transformed | p < 0.01 | Tukey's HSD: p<0.05 |
| | ANOVA; d.f.=3,99, F= 4.25 | $^{15}N$ treatment | | p < 0.01 | |
| | ANOVA; d.f.=3,99, F= 1.54 | Interaction | | p > 0.05 | |
| **S2c** | ANOVA; d.f.=1,51, F=15.5000 | Fungal inoculation | N/A | p < 0.001 | Tukey's HSD: p<0.05 |
| | ANOVA; d.f.=2,51, F=15.7688 | N concentration | | p < 0.001 | |
| | ANOVA; d.f.=2,51, F=0.3617 | Interaction | | p > 0.05 | |

#### Table S2 T-tests

**Table S2 Pairwise T-test statistics for hyphal growth at each time point (1-7; Fig. 4c)**

| Time point | Treatment 1 | Treatment 2 | $T_{13}$ | p value | Significance code |
|---|---|---|---|---|---|
| **1** | 187.4 μg.gN$^{-1}$ | 93.7 μg.gN$^{-1}$ | 0.2919211 | $p > 0.05$ | ns |
| | 187.4 μg.gN$^{-1}$ | 25 μg.gN$^{-1}$ | -5.4530972 | $p < 0.001$ | *** |
| | 93.7 μg.gN$^{-1}$ | 25 μg.gN$^{-1}$ | -4.7877685 | $p < 0.01$ | ** |
| **2** | 187.4 μg.gN$^{-1}$ | 93.7 μg.gN$^{-1}$ | -1.9652583 | $p > 0.05$ | ns |
| | 187.4 μg.gN$^{-1}$ | 25 μg.gN$^{-1}$ | -5.0535088 | $p < 0.001$ | *** |
| | 93.7 μg.gN$^{-1}$ | 25 μg.gN$^{-1}$ | -4.8184044 | $p < 0.01$ | ** |
| **3** | 187.4 μg.gN$^{-1}$ | 93.7 μg.gN$^{-1}$ | -2.164201 | $p > 0.05$ | ns |
| | 187.4 μg.gN$^{-1}$ | 25 μg.gN$^{-1}$ | -5.4294239 | $p < 0.001$ | *** |
| | 93.7 μg.gN$^{-1}$ | 25 μg.gN$^{-1}$ | -4.3429743 | $p < 0.01$ | ** |
| **4** | 187.4 μg.gN$^{-1}$ | 93.7 μg.gN$^{-1}$ | -1.4703968 | $p > 0.05$ | ns |
| | 187.4 μg.gN$^{-1}$ | 25 μg.gN$^{-1}$ | -4.2385481 | $p < 0.01$ | ** |
| | 93.7 μg.gN$^{-1}$ | 25 μg.gN$^{-1}$ | -3.1249716 | $p < 0.05$ | * |
| **5** | 187.4 μg.gN$^{-1}$ | 93.7 μg.gN$^{-1}$ | -3.4933944 | $p < 0.05$ | * |
| | 187.4 μg.gN$^{-1}$ | 25 μg.gN$^{-1}$ | -4.6848381 | $p < 0.01$ | ** |
| | 93.7 μg.gN$^{-1}$ | 25 μg.gN$^{-1}$ | -1.543355 | $p > 0.05$ | ns |
| **6** | 187.4 μg.gN$^{-1}$ | 93.7 μg.gN$^{-1}$ | -4.8014318 | $p < 0.01$ | ** |
| | 187.4 μg.gN$^{-1}$ | 25 μg.gN$^{-1}$ | -7.8208239 | $p < 0.001$ | *** |
| | 93.7 μg.gN$^{-1}$ | 25 μg.gN$^{-1}$ | -2.366826 | $p > 0.05$ | ns |
| **7** | 187.4 μg.gN$^{-1}$ | 93.7 μg.gN$^{-1}$ | -4.6667448 | $p < 0.01$ | ** |
| | 187.4 μg.gN$^{-1}$ | 25 μg.gN$^{-1}$ | -6.173636 | $p < 0.001$ | *** |
| | 93.7 μg.gN$^{-1}$ | 25 μg.gN$^{-1}$ | -1.2448251 | $p > 0.05$ | ns |

Table S3 Nutrient composition of media in 'N concentration' experiment

| Compound | M.Wt | g.L (High N) | g.L (Medium N) | g.L (Low N) |
|---|---|---|---|---|
| **Calcium chloride** | 110.98 | 0.05662 | 0.05662 | 0.05662 |
| **Magnesium sulfate.7$H_2O$** | 246.47 | 0.12538 | 0.12538 | 0.12538 |
| **Manganese sulphate.$H_2O$** | 151.00 | 0.00500 | 0.00500 | 0.00500 |
| **Ferrous sulphate.7$H_2O$** | 278.01 | 0.01390 | 0.01390 | 0.01390 |
| **EDTA disodium salt.2$H_2O$** | 372.24 | 0.01865 | 0.01865 | 0.01865 |
| **Zinc sulphate.7$H_2O$** | 287.60 | 0.00100 | 0.00100 | 0.00100 |
| **Boric acid** | 61.84 | 0.00150 | 0.00150 | 0.00150 |
| **Potassium iodide** | 166.00 | 0.00038 | 0.00038 | 0.00038 |
| **Molybdic acid (sodium salt).2$H_2O$** | 241.95 | 0.00013 | 0.00013 | 0.00013 |
| **Copper sulphate.5$H_2O$** | 249.69 | 0.00001 | 0.00001 | 0.00001 |
| **Cobalt chloride.6$H_2O$** | 237.93 | 0.00001 | 0.00001 | 0.00001 |
| **myo-Inositol** | 180.16 | 0.05000 | 0.05000 | 0.05000 |
| **Nictotinic acid (free acid)** | 123.11 | 0.00050 | 0.00050 | 0.00050 |
| **Pyridoxine • HCl** | 205.64 | 0.00050 | 0.00050 | 0.00050 |
| **Thiamine • HCl** | 337.27 | 0.00500 | 0.00500 | 0.00500 |
| **Sodium phosphate monobasic** | 119.98 | 0.06525 | 0.06525 | 0.06525 |
| **Potassium nitrate ($KNO_3$)** | 101.10 | 0.166250 | 0.62500 | 1.25 |
| **Ammonium sulphate ($(NH4)_2$ SO4)** | 132.14 | 0.008911 | 0.03350 | 0.067 |
